# Branched actin networks mediate macrophage-dependent host microbiota homeostasis

**DOI:** 10.1101/2024.07.18.604111

**Authors:** Luiz Ricardo C. Vasconcellos, Shaina Chor Mei Huang, Alejandro Suarez-Bonnet, Simon Priestnall, Probir Chakravarty, Sunita Varsani-Brown, Matthew L. Winder, Kathleen Shah, Naoko Kogata, Brigitta Stockinger, Michael Way

## Abstract

Assembly of branched actin networks, driven by the Arp2/3 complex are essential for the function and integrity of the immune system. Patients with loss-of-function mutations in the ARPC5 subunit of the Arp2/3 complex develop inflammation and immunodeficiency after birth, leading to early mortality. However, the mechanistic basis for these phenotypes remains obscure. Here we demonstrate that loss of Arpc5 in the murine hematopoietic system, but not the corresponding Arpc5l isoform causes early-onset intestinal inflammation after weaning. This condition is initiated by microbiota breaching the ileal mucosa, leading to local and systemic inflammation. Macrophage and neutrophils infiltrate into the ileum, but in the absence of Arpc5 fail to restrict microbial invasion. Loss of Arpc5 compromises the ability of macrophages to phagocytose and kill intra-cellular bacteria. Our results underscore the indispensable role of Arpc5, but not Arpc5l containing Arp2/3 complexes in mononuclear phagocytes function and host-microbiota homeostasis.

**One-Sentence Summary:** Arpc5 containing Arp2/3 complexes are essential for host-microbiota homeostasis

## Introduction

Mutations in genes encoding proteins involved in the regulation and organization of the actin cytoskeleton frequently result in primary immunodeficiency (*1*). These so-called actinopathies have a wide spectrum of severity involving the innate and/or the adaptive arm of the immune system (*2*). The most well-known actinopathy, Wiskott-Aldrich Syndrome (*3*), is caused by mutations in WASP, a hematopoietic specific protein that activates the Arp2/3 complex to nucleate branched actin filament networks (*4, 5*). A functioning immune system depends on these networks to drive cell migration, phagocytosis and immunological synapse assembly (*6, 7*). The Arp2/3 complex consists of seven evolutionarily conserved subunits, two Actin Related Proteins: Arp2, Arp3 and ArpC1-ArpC5 (*8, 9*). However, in mammals, three of the subunits have two different isoforms (Arp3/Arp3B, ARPC1A/ARPC1B and ARPC5/ARPC5L), each of which confers different properties to the Arp2/3 complex (*10–12*). The importance of these subunit isoforms is underscored by the observation that mutations in human ARPC1B result in severe immunodeficiency as well as impaired cytotoxic T lymphocyte maintenance and activity (*13–16*). More recently, loss of function mutations in ARPC5 have also been reported to result in gastrointestinal conditions, increased susceptibility to infection leading to sepsis and early death (*17, 18*). The mechanistic basis for these clinical observations in patients with ARPC5 mutations remains unknown. We therefore examined the impact of the loss of Arpc5 in the murine hematopoietic compartment to determine the basis for these immune phenotypes.

### Loss of Arpc5 in immune cells leads to spontaneous enteritis

As ubiquitous loss of Arpc5 in mice is embryonic lethal (*18*), we used the Vav1-iCre driver (*19*), to delete Arpc5 in the hematopoietic compartment (fig. S1A). At 8-15 weeks of age, *Arpc5*^fl/fl_Vav1Cre+^ animals (hereafter referred to as C5^ΔVav^) had reduced weight gain and increased levels of faecal lipocalin-2, a marker of intestinal inflammation (*20*), compared to littermate controls (C5^fl/fl^ and C5^HetVav^) (Fig.1A, fig. S1B). Consistent with this, adult C5^ΔVav^ had intestinal inflammation with increased infiltration of monocytes/macrophages and neutrophils, and loss of structural integrity, which was not seen in control mice (Fig.1B-C, fig. S1C-D). Single cell RNAseq analysis revealed that only monocytes/macrophages and neutrophils increase in the intestine of C5^ΔVav^ animals (Fig.1D). The inflammation and damage was restricted to the ileum of the small intestine (Fig. S1D-E). The mesenteric (mLNs) but not the inguinal lymph nodes (iLNs) were enlarged (Fig.1E), indicative of a local immune response in the intestine. The increase in size of mLNs was driven by an expansion of lymphocytes and significant recruitment of neutrophils and macrophages (fig. S1F). Importantly, no inflammation or intestinal damage were observed in age matched *Arpc5l*^fl/fl_Vav1Cre+^ (C5L^ΔVav^) animals (fig. S1G-H). Our observations demonstrate that Arpc5 but not the alternative Arpc5l isoform is essential for maintenance of intestinal homeostasis by the immune system.

**Figure 1.**
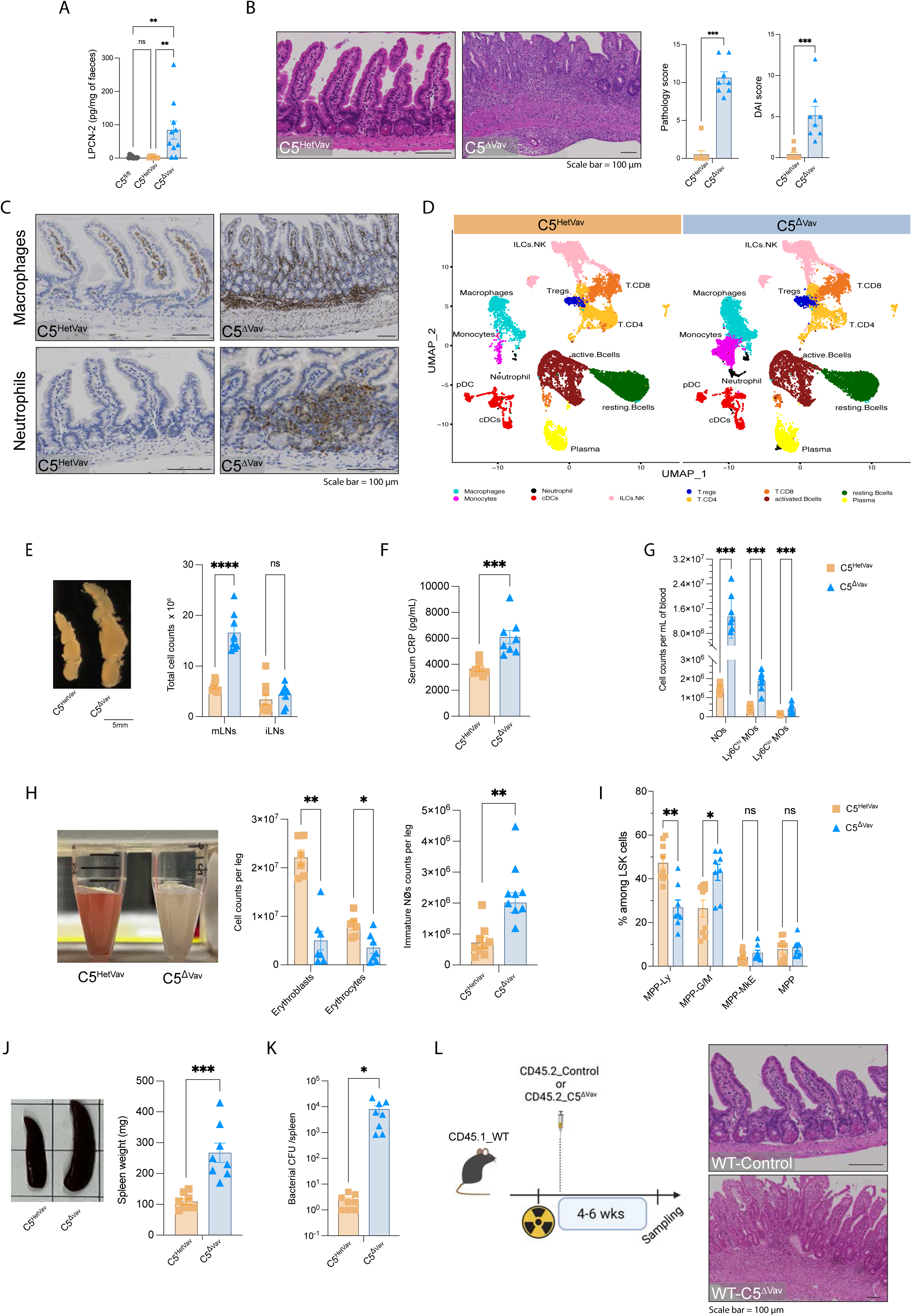
Arpc5 loss in immune cells leads to intestinal damage and systemic inflammation. (**A**) Lipocalin-2 (LPCN-2) faecal levels in the indicated mice. (**B**) H&E images of intestines (left) together with quantification of the histopathology score (middle) and disease activity index (DAI) (right). (**C**) Immunohistochemistry reveals the increased prevalence of macrophages (top) and neutrophils (bottom) in ileal sections in mice lacking Arpc5. **(D)** Uniform manifold approximation and projection (UMAP) of the integrated scRNAseq from C5^HetVav^ or C5^ΔVav^ ileum lamina propria. Cell clusters are colored by the assigned cell cluster indicated in the figure. (**E**) Loss of Arpc5 increases the size and cell counts of Mesenteric lymph nodes (mLNs) but not the inguinal lymph node (iLN). (**F-G**). Mice lacking Arpc5 have increased levels C-reactive protein (CRP) as well as phagocytes in their blood. **(H)** Representative image of bone marrow together with the quantification of the counts of erythroid lineage and immature neutrophil (NØ) in the bone marrow of the indicated mice. (**I**) Quantification of the % of multipotent progenitors (MPP) among the lineage^−^Sca1^+^c-Kit^+^ (LSK) cells. MPP-Ly: lymphoid-biased MPPs; MPP-G/M: myeloid-biased MPPs; MPP-MkE: megakaryocyte and erythroid-biased MPPs; MPP: unbiased MPPs. (**J**) Representative image and weight of spleen in the indicated mice. (**K**) Bacterial colony forming units (CFU) in C5^HetVav^ or C5^ΔVav^ mice spleens. (**L**) Schematic of the bone marrow transplantation protocol: CD45.1_wild type mice were lethally irradiated and reconstituted with either CD45.2_controls (WT-Control) or CD45.2_C5^ΔVav^ (WT-C5^ΔVav^) bone marrow (left). H&E images of intestines of animals which received wild-type (WT-control) or C5^ΔVav^ (WT-C5^ΔVav^) bone marrow transplantation (right). Animals assessed in in L were 8-15 week old mice n=7 of two independent experiments. For all the other panels, data shown of at least three independent experiments, n=8. All data in this figure are represented as means ± SEM. Statistical analysis was performed using Kruskal-Wallis test in **A**, Mann-Whitney test in **B E, F**, **H, J, K,** or multiple Mann-Whitney test in **G** and **I**. Significant *p* values are indicated on the graphs. C5^fl/fl^ (arpc5^fl/fl^), C5^HetVav^ (arpc5^HetVav^) and C5^ΔVav^ (arpc5^ΔVav^). Scale bars = 100µm. ns = not significant. * = p-value < 0.05. ** = p-value < 0.01. *** = p-value < 0.001 and **** = p-value < 0.0001.

### Arpc5 deficiency leads to systemic inflammation

Assessing the severity of inflammation, we found that C5^ΔVav^ mice had increased serum levels of C-reactive protein (CRP), a marker of inflammation (Fig.1F). The number of neutrophils and monocytes in the blood of C5^ΔVav^ mice increased, while only CD4 T cells decreased (Fig.1G and fig. S1I). Similar changes in cell numbers in the peripheral blood were reported in patients lacking ARPC5 (*17*). C5^ΔVav^ mice also had pale bone marrow due to a reduction in erythroid lineages and an increase in Ly6G^lo^ immature neutrophils (Fig.1H), as observed in mice with sepsis (*21*). Consistent with the substantial increase in myeloid cell counts, the proportion of myeloid-biased multipotent progenitors (MPP^G/M^) increased while lymphoid-biased ones (MPP^Ly^) decreased (Fig. 1I). Compared to controls, C5^ΔVav^ mice also had splenomegaly with increased numbers of neutrophils, macrophages, and dendritic cells (DCs), and reduced counts of natural killer cells but no changes in B and T cells (Fig.1J, fig. S1J). Indicative of microbial dissemination, the spleens of C5^ΔVav^ animals contained significant numbers of bacteria compared to the controls (Fig.1K). To determine whether the phenotype observed is cell intrinsic, we performed adoptive transfer of Arpc5 knockout (KO) bone marrow into lethally irradiated wild-type mice (Fig.1L). These animals developed the pathology observed in C5^ΔVav^ mice, confirming that the phenotype is hematopoietic compartment intrinsic (Fig.1L and fig. S1K).

### The microbiota drives the inflammatory response in C5^ΔVav^ animals

At weaning there is a significant change in the intestinal microbiota that profoundly impacts mucosal homeostasis (*22*). To investigate the onset of pathologies induced by the absence of Arpc5, we assessed animals immediately after weaning (4 week old). We found there was no significant increase in faecal lipocalin-2 levels, mLN enlargement or intestinal pathology in 4 week old C5^ΔVav^ animals compared to control littermates (Fig.2A, fig. S2A). Longitudinal RNAseq analysis of intestinal macrophages from animals at 4, 6 and 8 weeks revealed that differentially expressed genes (DEG) pathways were mainly associated with inflammation, but only in adult mice (8 week old, fig. S2B). The elevated numbers of macrophages and DCs in 4 week old C5^ΔVav^ mLNs (fig. S2C), however, suggests there is a progressive onset of inflammation after weaning. To explore whether local and systemic inflammation are triggered by commensal colonization after weaning, we treated 4 week old C5^ΔVav^ animals with an antibiotic cocktail prior to development of enteritis (Fig. 2B). This treatment prevented the development of enteritis, infiltration of macrophages and neutrophils into the ileum as well as subsequent systemic inflammation (Fig. 2C-E). Antibiotic treatment of C5^ΔVav^ mice also suppressed splenomegaly (Fig. 2F fig.S2D), enlargement of mLNs and maintained normal cell populations in the bone marrow (Fig.2G, fig.S2E). Analysis of the intestinal microbiome revealed that the loss of Arpc5 in the immune system also changed the composition of the microbiota significantly (Fig. 2H and fig.S2F). Interestingly, even before the intestinal damage was evident, there was an increase and decrease in clostridia and bacteroidia classes respectively in 4 week old C5^ΔVav^ animals (Fig. 2A and H). This change in microbiota composition was not seen before weaning (2week) (Fig. 2H and fig.S2F). Our observations demonstrate the onset of intestinal inflammation is progressive and driven by the microbiota.

**Figure 2.**
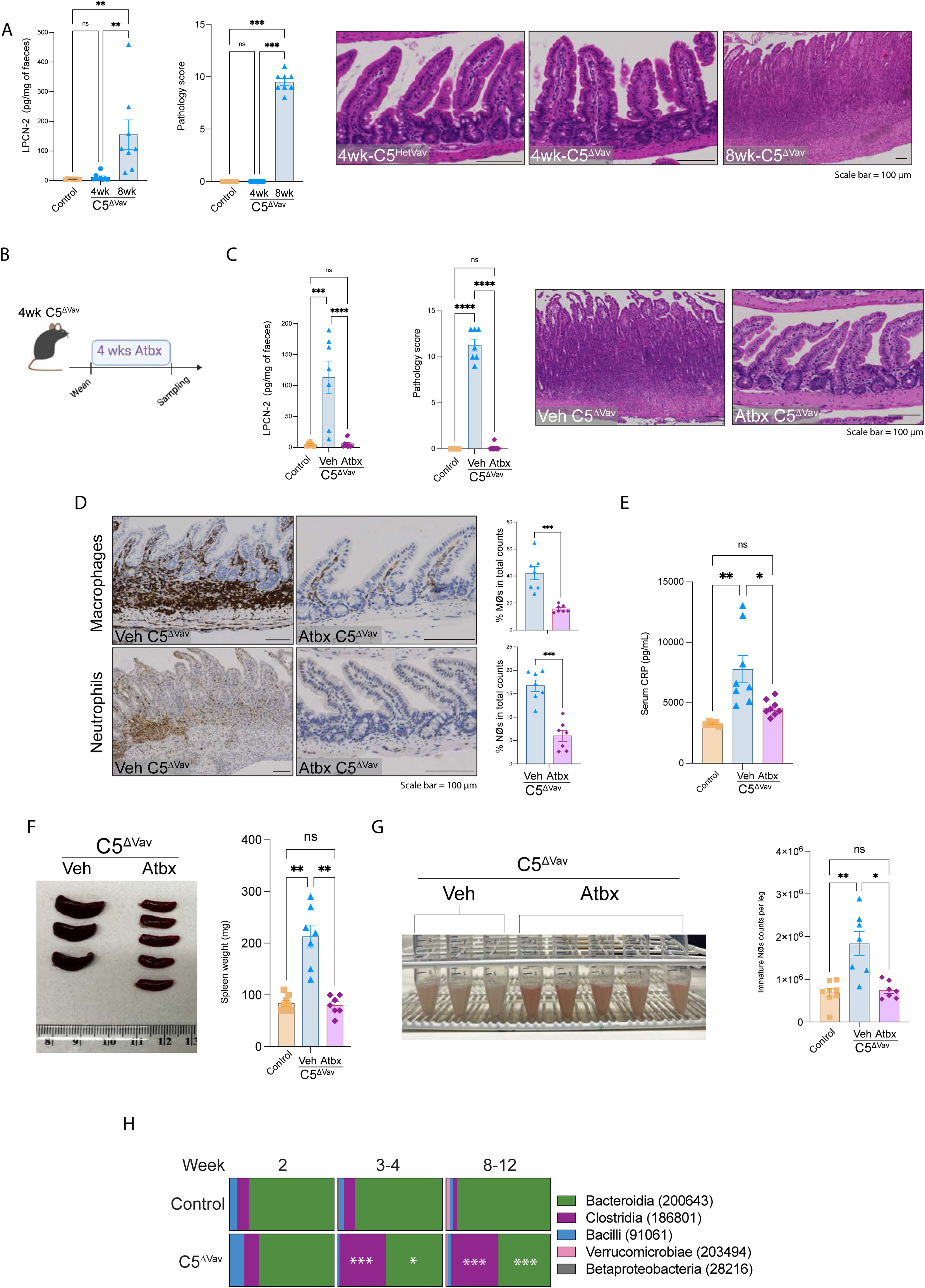
The intestinal dyshomeostasis upon Arpc5 loss is microbiota dependent. (**A**) Faecal lipocalin-2 (LPCN-2) levels (left), quantification of the histopathology score (middle) and H&E images of the ileum of 4 week old (wk) C5^HetVav^ or 4 and 8wk C5^ΔVav^ animals (right). (**B**) Experimental scheme of antibiotic treatment; 4wk animals were treated with antibiotic cocktail (Atbx) or vehicle (Veh) for 4 weeks and then sampled for analysis. (**C**) Quantification of LPCN-2 levels in faeces (left), and histopathology score (middle) together with H&E image of the ileum in the indicated mice (right). (**D**) Representative immunohistochemistry images and quantification showing Atbx treatment reduces macrophages (top) and neutrophils (bottom) presence in ileal sections in C5^ΔVav^ mice. **(E**) Levels of C-reactive protein (CRP) in the blood of animals with and without antibiotic treatment. **(F**) Representative image and weight of spleens in C5^ΔVav^ animals with and without antibiotics. (**G**) Representative image of bone marrow together with the quantification of numbers of immature neutrophil (NØ) in the bone marrow of mice with and without antibiotic treatment. (**H**) Five most prevalent classes of bacteria found in the intestinal microbiome of control (C5^HetVav^) or C5^ΔVav^ animals at the indicated age. Data shown as means ± SEM of at least three independent experiments, n=7, except for **H** where n=8. Statistical analysis was performed using Kruskal-Wallis tests in **A**, **C**, **E**, **F** and **G,** or Mann-Whitney test in **D;** Multiple Mann-Whitney test was applied in **H**. Significant *p* values are indicated on the graphs. Controls used were Arpc5^HetVav^ and C5^ΔVav^ (Arpc5^ΔVav^). Scale bars = 100µm. ns = not significant. * = p-value < 0.05. ** = p-value < 0.01. *** = p-value < 0.001 and **** = p-value < 0.0001.

### Arpc5 is essential for intestinal homeostasis

We next sought to determine whether the phenotype seen in C5^ΔVav^ mice was driven by a deficiency in innate and/or adaptive immune cells. The scRNAseq analysis of immune cells in the intestine demonstrated that innate immune cells increased in C5^ΔVav^ mice (monocytes/macrophages and neutrophils, fig. S3A). Furthermore, DEG pathway analysis revealed increased transcription of genes associated with actin cytoskeleton dynamics, inflammation/innate immunity, extracellular matrix remodelling and oxidative stress (fig. S3B). In contrast, there was no significant difference in cell numbers in the adaptive compartment or impact in polyclonal activation in C5^ΔVav^ T cells isolated from mLNs (fig. S3A and S3C). Similarly, no defect in T cell response was reported in ARPC5 mutant patients (*17*). Since an interaction between monocytes/macrophages and Tregs in the intestine is essential for maintaining tolerance (*23*), we assessed the cell-to-cell crosstalk in our scRNAseq dataset using CellChat (*24*). We found there were increased differential interactions of monocytes/macrophages with themselves as well as with Tregs and other immune cells in the absence of Arpc5 (fig. S3D). Furthermore, it is striking that Tregs do not produce interleukin-10 (IL-10) in C5^ΔVav^ animals (fig. S3E), given it is required for the tolerogenic response (*25*). Consistent with the lack of IL-10 production, Arpc5 knockout Tregs failed to maintain intestinal homeostasis in a lymphocyte transfer model (fig. S3F).

To investigate the contribution of the adaptive immune system in the intestinal phenotype induced by the absence of Arpc5, we crossed C5^ΔVav^ mice on to a *Rag^−/−^* background (C5^ΔVav/RagKO^), which lack T and B cells. C5^ΔVav/RagKO^ mice had an even more pronounced phenotype, with all animals reaching an humane endpoint around 6 weeks of life due to intestinal and systemic inflammation (Fig.3A-B and fig. S4A). Interestingly, in these C5^ΔVav/RagKO^ mice the intestinal damage was now also observed in the cecum (fig.S4A). In addition, there were similar changes in the microbiota of C5^ΔVav/RagKO^ animals as that seen in the C5^ΔVav^ mice (fig.S4B). Consistent with this, adoptive transfer of C5^ΔVav/RagKO^ bone marrow into lethally irradiated wild-type mice also transferred the pathological phenotype (Fig.3C-D and fig. S4C-D). This confirms that the phenotypes seen in C5^ΔVav^ mice are driven by deficiencies in innate immunity.

**Figure 3.**
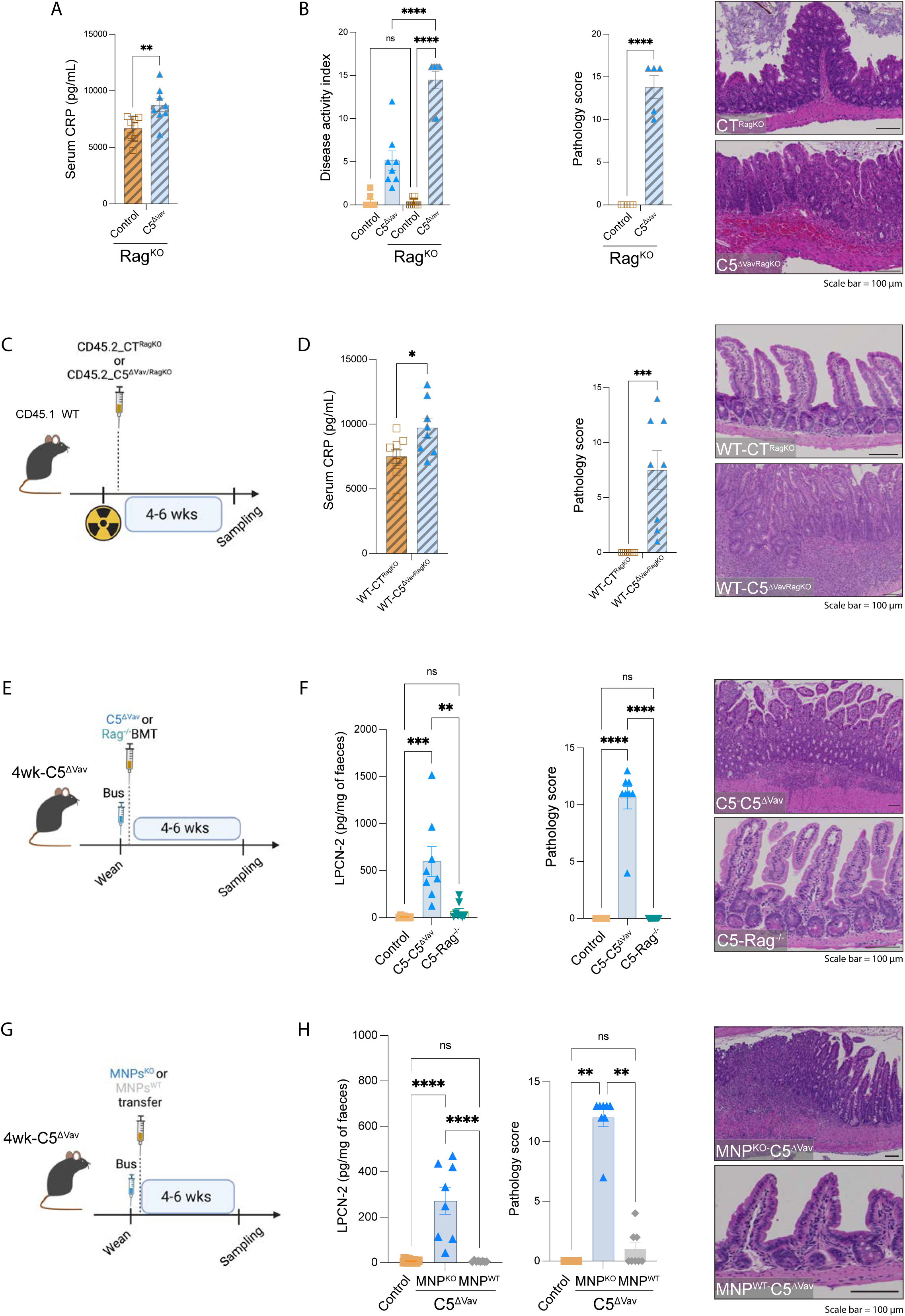
Arpc5 is essential for effective innate immunity. (**A**) Levels of C-reactive protein (CRP) in the serum and (**B**) quantification of disease activity index (DAI including values also shown in Fig.1C, left) and histopathology score together with representative H&E images of cecum of control^RagKO^ or C5^ΔVavRagKO^ mice (right). (**C**) Schematic of the bone marrow transplantation protocol: CD45.1_wildtype mice were subjected to lethal irradiation and reconstituted with either CD45.2_control^RagKO^ (WT-CT^RagKO^) or CD45.2_C5^ΔVavRagKO^ (WT-C5^ΔVavRagKO^) bone marrow. (**D**) Levels of CRP (left) and quantification of the histopathology score in irradiated mice together with representative H&E images of ileum (right). (**E**) Scheme for Rag^−/−^ adoptive cell transfer after busulfan treatment: 4 week old C5^ΔVav^ mice were treated with busulfan and injected with either WT (C5-Rag^−/−^) or C5^ΔVav^ (C5-C5^ΔVav^) innate cells. (**F**) Lipocalin-2 (LPCN-2) faecal levels (left) and quantification of the histopathology score together with representative H&E images of the ileum of busulfan-treated mice (right). (**G**) Scheme for mononuclear phagocytes (MNPs) adoptive transfer into 4 week old C5^ΔVav^ busulfan depleted mice: animals were transplanted with either WT (MNP^WT^) or C5^ΔVav^ (MNP^KO^) cells. (**H**) LPCN-2 faecal levels (left) and quantification of the histopathology score together with representative H&E images of the ileum (right) of indicated animals. Data shown as means ± SEM of at least three independent experiments, n=8 except as indicated in (H). Statistical analysis was performed using Mann-Whitney test in **A**, **B** (right) and **D;** multiple Mann-Whitney test in **B** (left) and Kruskal-Wallis tests in **F** and **H**. Significant *p* values are indicated on the graphs. Controls used were Arpc5^HetVav/RagKO^. C5^ΔVavRagKO^ (Arpc5^ΔVav/RagKO^). Scale bars = 100µm. ns = not significant. * = p-value < 0.05. ** = p-value < 0.01. *** = p-value < 0.001 and **** = p-value < 0.0001.

The current treatment for Wiskott-Aldrich Syndrome involves transplantation of hematopoietic stem/progenitor cells (*26, 27*). We decided to use a similar strategy, performing adoptive transfer of innate immune cells derived from *Rag*^−/−^ bone marrow into 4 week old C5^ΔVav^ animals after myeloablation (Fig.3E). We found that C5^ΔVav^ animals reconstituted with wild-type cells (C5-Rag^KO^) had no signs of inflammation or the pathologies seen in C5^ΔVav^ mice (Fig. 3F, fig. S4E-F).

Mononuclear phagocytes (MNPs) play a key role in safeguarding intestinal homeostasis by clearing microbes that breach the ileal mucosa (*28–31*). Given this, we performed adoptive transfer of CD115^+^ MNPs from wildtype or C5^ΔVav^ bone marrow (MNPs^WT^-C5^ΔVav^ and MNPs^KO^-C5^ΔVav^) into myeloablated 4 week old C5^ΔVav^animals (Fig. 3G). Strikingly, MNPs^WT^-C5^ΔVav^ animals developed no enteritis or splenomegaly and have comparable levels of circulating CRP and faecal lipocalin-2 to controls (Fig. 3H and fig. S4G). Similarly, wildtype MNPs transfer after peripheral depletion with clodronate liposomes ameliorated the pathology in C5^ΔVav^ animals, although inflammation was still present (fig. S4H). Our observations confirm the importance of innate immune cells and more specifically macrophages for the inflammatory phenotypes observed in the absence of Arpc5.

### Arpc5 is required for efficient phagocytosis and bacterial killing

Arpc5 deficient bone marrow-derived macrophages (BMDMs) have striking differences in cell morphology, being less spread and more elongated than wild-type counterparts (Fig.4A, fig. S5A). These differences indicate that loss of Arpc5 impacts the organisations of the actin cytoskeleton as reported in other cell types (*32*) (*33*). Our histological analysis, however, indicated that macrophages without Arpc5 readily infiltrate into the small intestine (Fig 1C), suggesting that defects in cell migration are not a major contributing factor in the phenotype of C5^ΔVav^ mice. Nevertheless, it was noticeable that BMDMs lacking Arpc5 had reduced levels of F-actin indicative of altered actin dynamics (Fig.4B, fig. S5B). It is well established that microbial internalization by phagocytes is actin dependent and defects in this process may contribute to inflammatory bowel disease (IBD) (*32, 34*). Consistent with this, we found that BMDMs lacking Arpc5 were less efficient in phagocytosing *E. coli* than controls and were defective in actin filament assembly (Fig.4B-C and fig. S5C-D). Live cell imaging of BMDMs expressing GFP-LifeAct revealed that the absence of Arpc5 resulted in a loss of actin rich ruffles required for bacterial phagocytosis leading to a reduction in their uptake (Fig 4D and E; movie 1 and 2). Furthermore, in contrast to wild-type BMDMs, the loss of Arpc5 also led to reduced bacterial killing as the number of intracellular bacteria remained constant (Fig.4F). Consistent with their defective bactericidal activity, infected BMDMs lacking Arpc5 were also more prone to die (Fig.4G). Our analyses support the notion that Arpc5 deficient macrophages are defective in phagocytosing and killing microbes that breach the intestinal barrier.

**Figure 4.**
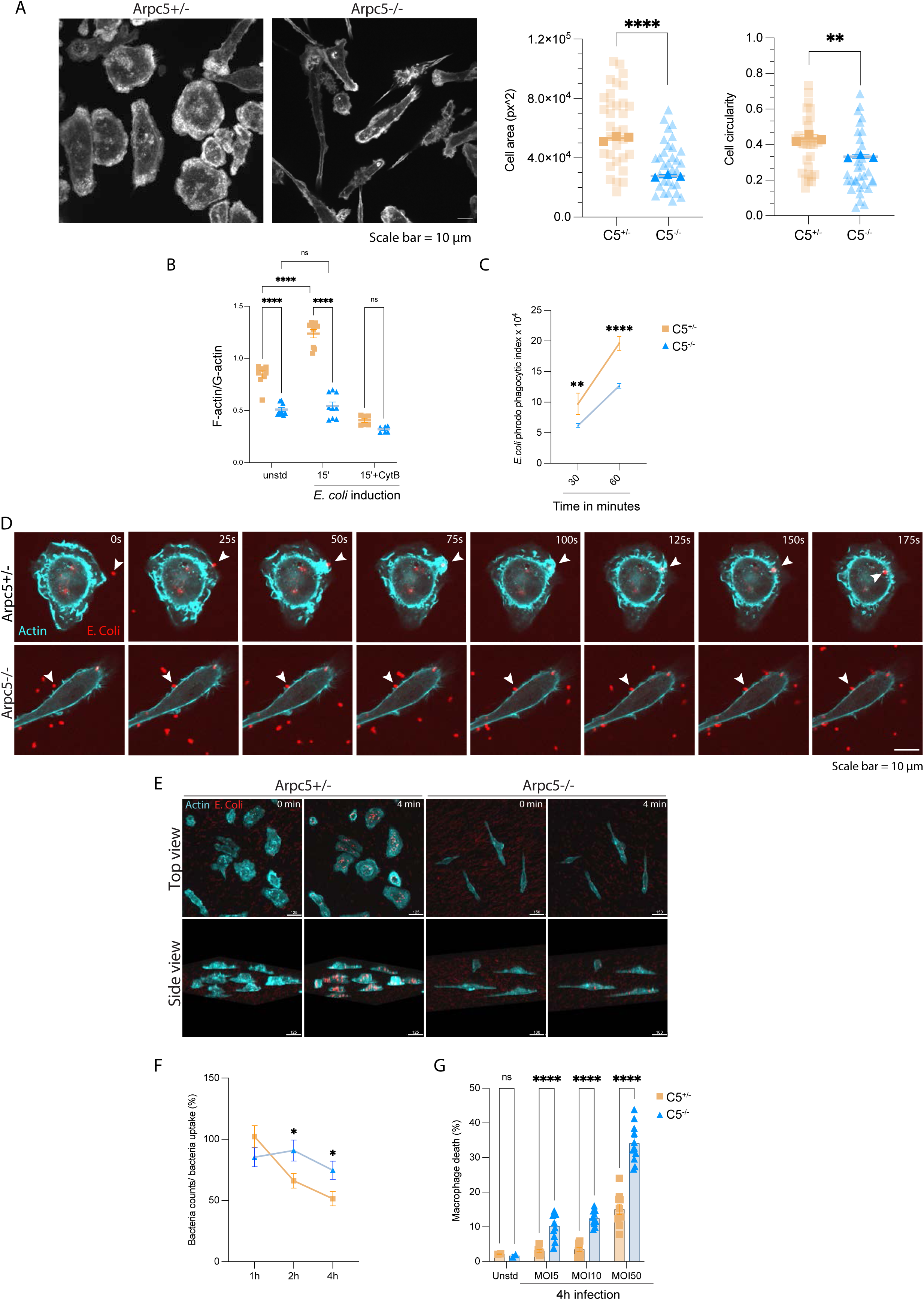
ArpC5-mediated actin remodeling is key to macrophage biology. **(A)** Representative images of the actin cytosksleton in bone marrow derived macrophages (BMDMs) with or without Arpc5 together with quantification of their cell area and circularity. (**B**) Ratio of F-actin/G-actin in resting or *E.coli*-activated BMDMs incubated with vehicle or pretreated with 10µM Cytochalasin D for 30min. (**C**) Quantification of *E. coli* phrodo particle uptake by BMDMs. (**D**) Representative images showing the actin cytoskeleton dynamics (Cyan) visualized with GFP-Lifeact in wildtype and Arpc5^−/−^ BMDMs during *E. coli* (Red) phagocytosis. The time in each panel is seconds. (**E**) Representative images showing top and side views (3D representation) of *E. coli* (Red) in BMDMs. Scale bar (µm) are indicated in each panel. (**F**) The quantification of the percentage of internalized bacteria by BMDMs. (**G**) *E. coli* infection induced BMDM cell death. Data shown as means ± SEM of at least three independent experiments. Statistical analysis was performed using unpaired *t* test in **A**; two-way analysis of variance (ANOVA) in **B, C** and **F** and multiple Mann-Whitney test in **G**. Significant *p* values are indicated on the graphs. C5^+/−^ = Arpc5^+/−^, C5^−/−^ = Arpc5^−/−^. Scale bars = 10µm. ns = not significant. * = p-value < 0.05. ** = p-value < 0.01. and **** = p-value < 0.0001.

Our analysis has uncovered that intestinal homeostasis depends on Arpc5, but not Arpc5l containing Arp2/3 complexes nucleating branched actin networks in innate immune cells. While the inflammatory effect observed in Arpc5 mutants does not require the adaptive immune system, it is likely to be exacerbated by defective Tregs that cannot produce IL-10. Interestingly, Arpc5 dependent innate immunity also regulates the composition of the intestinal microbiota prior to the onset of intestinal inflammation. Furtherwork will be required to understand how Arpc5-dependent actin polymerization in innate immune cells regulates the composition of the microbiota. Our observations, however, demonstrate that the Arpc5 deficiency pathology is not dependent on the ability of macrophages to migrate to the intestine but rather their inability to perform phagocytosis and microbial killing. Interestingly, the Arp2/3 complex is also not required for dendritic cells to migrate to the intestinal lamina propria (*35*), but a functional Arp2/3 complex is required for macrophage phagocytosis (*36–38*).

In summary, our study now explains why patients lacking ARPC5 develop gastrointestinal complications, immunodeficiency and are prone to fatal sepsis. We propose that mutations in ARPC5 may contibute to IBD and should be considered in patients with mutations in other candidate genes, such as ARPC1B and WAS (*33, 39*). Finally, our results suggest that bone marrow transplantation would revert the immunodeficiency of patients with ARPC5 loss of function mutations.

## Supporting information

Combined revised supplemental figures

## Acknowledgments

We thank Patrick Costello, Chrysante Iliakis, and all staff in the Crick Scientific Technology Platforms: Genetic Modification Service, Biological Research Facility, Experimental Histopathology, Cell Science, Bioinfomatics and Biostatistics, Advanced Light Microscopy, Genomics Science Technology and Flow Cytometry STPs for technical support. We thank particularly Hubert Slawinski, Christian Heafield and Marg Crawford, for their contributions to the single-cell capture experiment, library construction and sequencing. We also thank Rocco D’Antuono for his help with microscopy and 3D image analysis and Emanuele Libertini for the technical support for bulkRNAseq data analysis. We also thank James Lee, Jeremy Carlton, Caetano Reis e Sousa, Andreas Wack, Maximiliano Gutierrez and Venizelos Papayannopoulos for providing feedback on the manuscript as well as Richard Treisman and Andreas Wack for sharing reagents. Experimental schematics in the paper were created using BioRender. For the purpose of open access, the authors have applied a CC BY public copyright licence to any author accepted manuscript version arising from this submission.

## Funding

European Research Council (ERC) under the European Union’s Horizon 2020 research and innovation programme grant agreement No 810207 (MW). Francis Crick Institute, which receives its core funding from Cancer Research UK (CC2096) (MW) and (CC2016) (BS), the UK Medical Research Council (CC2096) (MW) and (CC2016) (BS), and the Wellcome Trust (CC2096) (MW) and (CC2016) (BS).

## Author contributions

Conceptualization: LRCV, MW, BS

Methodology: LRCV, SH, KS, NK, SB, MLW

Investigation: LRCV, SH, AS, SP, PC

Visualization: LRCV, SH, PC

Funding acquisition: MW

Project administration: MW, LRCV, SH, NK

Supervision: MW, BS

Writing – original draft: LRCV

Writing – review & editing: all authors.

## Competing interests

Authors declare that they have no competing interests.

## Data and materials availability

The scRNAseq and bulkRNAseq data associated with this study are publically available at the NCBI GEO repository with the identifiers GSE298330 and GSE298655, respectively. The raw and processed microbiome data associated with this study are publically available at One Codex (https://app.onecodex.com/projects/lvasconcellos). All data are available in the main text or the supplementary materials.

## Supplementary Materials

### Materials and Methods

#### Mice

In the present study we utilized sex and age-matched (4-15 weeks old) C57Bl/6 mice, from both genders, bred at the Francis Crick Institute (2019–2024) under pathogen-free conditions. Mice were genotyped before weaning by Transnetyx and randomly assigned to treatment groups and blinding strategies applied when possible. For adoptive transplantation, both *Ptprc^a^* (CD45.1) and *Ptprc^b^* (CD45.2) were used. The genetically modified animals *Arpc5*^Cre-Vav1^, *Arpc5*^Cre-Foxp3-YFP^*, Rag1*^−/−^, *Rag1*^−/−^/*Arpc5*^Cre-Vav1^, C57Bl/6^LifeAct-GFP^ and *Arpc5*^Cre-Vav1^/ ^LifeAct-GFP^ (*40*) were used in the present study. All in vivo experiments were performed following the Act 1986 for animal scientific procedures after the approval by the review board of The Francis Crick Institute and the Home Office (United Kingdom) under project licence PP0792028.

The conditional Arpc5 knockout mouse model has been previously reported (*18*). The Arpc5l conditional knockout was generated by the Crick Genetic Modification Service using a similar CRISPR-Cas9 approach in embryonic stem cells (ESCs). Briefly, the gRNA-1: 5’-CTCTGGATAGAGACAACAGA-3’ and gRNA-2: 5’-CATCCCCGATAGAGCTACAT-3’ were used to introduce the 5′ and 3′ loxP sites flanking exon 2 of the Arpc5l gene. Deletion of exon 2 results in a frameshift and an exogenous stop codon within exon 3. The Cas9-gRNA-Puro plasmids were generated by inserting the CRISPR-Cas9 target sequences into PX459 plasmid (Addgene plasmid #48139). The pMA donor plasmid contained a synthesized 1049 bp fragment corresponding to exon 2 flanked by loxP sites and 1 kb flanking homology arms (GeneArt, Thermo Fisher Scientific). Sequence-verified Cas9-gRNA-Puro plasmids and pMA donor plasmid were co-transfected into in-house C57BL6/N 6.0 ESC using Lipofectamine 2000 (Thermo Fisher Scientific) and selected with puromycin. Integration of 5′ and 3′ loxP sites were identified by reverse transcription quantitative polymerase chain reaction (RT-PCR) assays on genomic DNA using gene-specific probes designed by Transnetyx and the integrity of the loxP integrations was confirmed by PCR amplicons sequencing. ESC clones were microinjected into C57BL/6J mice blastocysts [B6(Cg)-Tyrc-2J/J; strain #000058, The Jackson Laboratory) and then transplanted into the uteri of pseudo-pregnant CD1 females. The resulting chimeras were crossed to albino C57BL/6J, and their offspring screened for the integration of the loxP sites by amplicon sequencing. Heterozygous floxed offspring were validated by sequencing and two founder strains maintained on a C57BL/6J background.

#### Disease activity index and histology

To assess the enteritis disease activity index (DAI), we followed a modified scoring system as described previously (*41*). Briefly, mice were evaluated using signs of clinical disease including weight loss, faecal consistency and presence of blood, and lack of movement. All parameters were assessed using a semi-quantitative grading scheme as follows: 0, no disease; 1, mild; 2, moderate; 3, marked; 4, severe.

Intestines (‘Swiss rolls’) and other organs were collected, fixed for 48h in 10% neutral buffered formalin which was subsequently replaced with 70% ethanol, embedded in paraffin wax and sectioned at 4µm. Sections were stained with H&E then microscopically examined by two certified veterinary pathologists blinded to the grouping and genotypes. Histopathological analyses considered the presence and extent of tissue inflammation and epithelial injury. All parameters were assessed in a semi-quantitative grading scheme for severity as follows: 0, no lesion; 1, minimal change; 2, mild change; 3, moderate change; 4, marked change; 5, severe.

Quantification of total leukocytes, macrophages and neutrophils immunohistochemistry was performed using anti-CD45, anti-F4/80 or anti-2B10, respectively, conjugated with a horseradish peroxidase (HRP) secondary antibody and using DAB chromogen. The slides were scanned with a Zeiss Axio Scan.Z1 annotated and processed with QuPath software (v0.4.3). To account for the prevalence of target cells, 10 different random regions (750μm^2^) were analyzed for positive labelling using hematoxylin as counterstaining. The values are presented as prevalence (%) of the Target cell/ Total cells.

#### Irradiated chimeras

CD45.1^+^ wild-type recipient mice at 10-13 weeks old were lethally irradiated with ^137^Cs-irradiated 2 × 6 Gy with a 3-hour interval. For reconstitution, 10 x 10^6^ bone marrow cells from donors (CD45.2^+^, as indicated in figures) were injected intravenously via tail vein into the recipients (CD45.1^+^). Bone marrow cells were collected from femur and tibia of donor mice and washed through a 70μm cell strainer. At 5-10 weeks later, the animals were euthanized, samples collected, and engraftment verified by flow cytometry. Distinction between donor and recipient cells was assessed by CD45.1 or CD45.2 expression.

#### Antibiotic treatment

Four week old *Arpc5*^Cre-Vav1^ (C5^ΔVav^) mice were treated continuously for 4 weeks in the drinking water with an antibiotic cocktail (Atbx) consisting of ampicilin (1mg/mL), gentamicin (1mg/mL), metronidazole (1mg/mL), neomycin (1mg/mL), vancomycin (500μg/mL) and sucralose-based artificial sweetener (100mg/mL, The Pantry) or artificial sweetener only (vehicle). Mice were euthanized, and samples collected after treatment.

#### Intestinal microbiome sequencing and analysis

Fresh faecal samples were collected from individual animals using the Microbiome collection kit (Transnetyx, Cordova, USA) and shipped to Transnetyx for DNA extraction using the Qiagen DNeasy 96 PowerSoil Pro QIAcube HT extraction kit (Qiagen, #47021). After DNA extraction and quality control, genomic DNA was converted into sequencing libraries using the KAPA HyperPlus library preparation protocol (Roche, 07962401001). Libraries were sequenced using the shotgun sequencing (a depth of 2 million 2×150 bp read pairs), using the Illumina NovaSeq instrument and protocol (Illumina, 20068232). Data were uploaded automatically onto One Codex analysis software (Wilmington, USA) and analyzed against the One Codex database of microbial reference genomes.

#### Busulfan chimeras

Four week old *Arpc5*^Cre-Vav1^ (C5^ΔVav^) mice were treated with 10mg/kg Busilvex (busulfan-Pierre Fabre) intraperitoneally for 2 consecutive days with a 24h interval. All donor bone marrow cells (10 x 10^6^) or 5 x 10^6^ CD115^+^ (isolated following the manufacturer’s instructions - Miltenyi Biotec, #130-096-354) in 100μl PBS were injected intravenously into the tail vein, 24h after the second busulfan dose (*42*). Bone marrow cells were collected from femur and tibia of donor mice and washed through a 70μm cell strainer. After adoptive cell transfer, animals were kept for 8-10 weeks before euthanasia and sample collection. CD45.1 and CD45.2 expression were used to differentiate between donor and recipient cells in transplanted animals.

#### In vivo macrophage depletion and adoptive cell transfer

Four week old *Arpc5*^Cre-Vav1^ (C5^ΔVav^) mice received intraperitoneal injection of 5μl/g clodronate_SUV_PEG liposomes (CP-SUV-P-005-005-Liposoma) and 48h later were adoptively transferred with 3 x 10^6^ CD115^+^ isolated from wild-type or C5^ΔVav^ mice bone marrows following the manufacturer’s instructions (Miltenyi Biotec, #130-096-354).

#### Immunoblot and Enzyme-linked immunosorbent assay (ELISA)

Cells and tissue extracts were lysed for 15 min on ice in RIPA buffer supplemented with protease and phosphatase inhibitors (Roche, catalogue number). Protein extracts were quantified, and similar amounts of total cell lysate separated by SDS-PAGE, followed by immunoblot analysis. Antibodies used for immunoblots: ArpC5 (Synaptic Systems, 305011), ArpC5L (Proteintech, 22025-a-AP), β-actin (Abcam, ab179467), anti-mouse IgG-IRDye® 800CW (LI-COR Biosciences, 926-32210) and anti-rabbit IgG-IRDye® 680RD (LI-COR Biosciences, 926-68071).

For LPCN-2 assessment, fresh faeces pellets were obtained and weighed prior to maceration in 300μl of PBS 0.1% triton X100 (Sigma) followed by ELISA detection. The levels of Lipocalin-2 (LPCN-2, DY1857-05) and C-reactive protein (CRP, MCRP00) were detected following the manufacturer’s instructions (R&D Systems).

#### Splenic Colony forming units (CFU)

Mouse spleens were dissociated in 500μl PBS 0.2% NP40 (Merck-492016) and centrifuged at 400x*g* for 3 min at room temperature to pellet debris. 100μl of the supernatant was plated on LB agar, incubated at 37°C for 24h and CFU determined.

#### Flow cytometry

Single cell suspension was preincubated with viability dye and FcgRIII/II (Fc block) before 30 min of incubation with fluorochrome-labelled antibodies or phalloidin. The stained cells were analyzed using BD LSR Fortessa cell analyser (BD Biosciences) and interpreted using Flowjo software (v.10.6.2). For intracellular staining cells were fixed and permeabilized using the Foxp3/Transcription Factor Staining Buffer Set (00-5523-00 ThermoFisher Scientific). FACs buffer (Thermo Fisher Scientific, 00-4222-26), Live/Dead^TM^ Fixable Near-IR Dead Cell Stain Kit (Thermo Fisher Scientific, L10119).

The following antibodies (supplied by Biolegend or the indicated company) were used in flow cytometry: CD16/32 Fcx (101320), B220-FITC (103205), B220-PerCP (103234), B220-APC (103212), CD2-FITC (100105), CD3-BV711 (100241), CD3-BUV395 (BD, 563565), CD4-BV605 (100548), CD4-BV711 (100557), CD8a-BV650 (100742), CD8a-BV737 (BD, 612759), CD8a-PE/Cy7 (100722), CD8b-BV786 (BD, 740952), CD11b-BV421 (101236), CD11b-BV711 (101242), CD11b-BV785 (101243), CD11c-AF488 (117311), CD11c-BV605 (117334), CD11c-BUV563 (BD, 749040), CD19-BV711 (115555), CD16/32-BV711 (101337), CD19-APC/Cy7 (BD, 557655), CD45-APC (103112), CD45-BUV395 (BD, 564279), CD45.1-AF700 (110724), CD45.2-BV785 (109839), CD48-PE (103406), CD64-PeCy7 (139314), CD71-PE (113807), CD103-AF488 (121408), CD103-BV605 (BD, 740355), CD117-APC (BD, 553356), CD135-BV421 (BD, 562898), CD150-BV605 (115927), F4/80 (Invitrogen, 17-4801-82), Foxp3-PE (126404), Foxp3-AF700 (126422), Gr-1-FITC (108406), Gr-1-BV711 (108443), I-A/I-E-AF488 (107616), I-A/I-E-AF700 (107622), Ly6C-BV785 (128041), Ly6C-APC (128016), Ly6G-FITC (BD, 551460), CD44-BV421 (103040), CD69-FITC (104506), NK1.1-BV711(108745), NK1.1-PE (BD, 557391), TCRab-BV421 (109230), TCRgd-APC (118116), TCRab-PE (eBioscience, 125961-82), Ter119-FITC (116206), Sca-1-PE/Cy7 (Invitrogen, 25-5981-82), AntiDNAse-I (Invitrogen, D12371) and CD45^rb^-AF647 (562848).

#### Leukocytes isolation from intestinal lamina propria

Leukocytes were isolated from mice intestines following an adapted protocol provided by Miltenyi Biotec (Lamina Propria Dissociation Kit, 130-097-410). Small intestines were isolated, and residual fat tissue and Peyer’s patches removed. The faeces were cleared and the tissue opened longitudinally in cold 10mM Hepes-PBS without Ca^+2^ and Mg^+2^ (PBSCM). Tissue was cut laterally into 0.5 cm length pieces and washed twice with predigestion solution (1mM DTT, 5mM EDTA, 5% FCS in PBSCM) for 20 min at 37°C with rotation. Samples were vortexed for 10 sec, filtered and incubated with rotation for 20 min at 37°C in PBSCM prior to being vortexed for 10 sec and filtered with a 100μm cell strainer. The strained samples were transferred to gentle MACS C Tube (Miltenyi Biotec, 130-093-237) containing pre-heated digestion solution (50μl Enzyme D, 50μl Enzyme R, 6.25μl Enzyme A) and digested in the program 37_m_LPDK_1 using the MACS Octo Dissociator with Heater (Miltenyi Biotec, 130-134-029). After digestion, samples were resuspended in FACS buffer (Thermo Fisher, 00-4222-26) and filtered through a 100μm cell strainer, cells counted and resuspended in appropriate volume for further assessment.

#### RNA-seq and analysis of ileum lamina propria macrophages

Macrophages were FACS sorted from ileum lamina propria of individual mice using a gating strategy to collect live cells, lineage^−^ (CD3/CD19/NK1.1/Ly6G) and CD45^+^CD11b^+^CD64^+^. RNA was extracted from samples using the QIAShredder and RNeasy Mini Kit with on-column DNase digestion, following manufacturer’s instructions (Qiagen, 74904). RNA-seq libraries were made using total RNA with KAPA RNA HyperPrep with RiboErase following manufacturer’s instructions (Roche, 08098131702). SMART-Seq libraries were pooled and sequenced on an Illumina NovaSeq 6000 (2 × 100 bp) (Illumina). Demultiplexed FASTQ files yielded a median 32.4 million paired reads per sample (IQR 4.7 M). All processing followed the nf-core/rnaseq pipeline v3.18.0 executed under Nextflow v24.04.2 with Singularity v3.6.4 on a high-performance cluster. Adapters and low-quality bases were removed with TrimGalore v0.6.10 (Cutadapt v4.9, FastQC v0.12.1). Post-QC, ≥ 98 % of bases per sample retained a Phred ≥ 30. Pipeline quality metrics from FastQC, Qualimap v2.3, RSeQC v5.0.2 and Samtools v1.21 were aggregated with MultiQC v1.18. Reference indices were built from the Mus musculus genome (GRCm38, Ensembl 95). Reads were mapped with STAR v2.7.11b in two-pass mode and simultaneously quantified in selective-alignment mode with Salmon v1.10.3. Transcript-level abundances were summarized to length-scaled gene-level counts with tximport v1.28.0; genes with < 10 raw counts in < 50 % of samples were excluded (edgeR filterByExpr, edgeR v3.42.0). Count matrices were imported into DESeq2 v1.38.3. Size-factors used the median-ratio method; dispersions were fitted to a log-normal prior. Variance-stabilised counts (vst) informed PCA, UMAP (umap v0.2.10.0) and distance heat-maps. Differential expression: a composite factor group (KOvsCTsgenotype × developmental stage) encoded six biological states. Global comparisons comprised: (i) within-genotype time-courses and (ii) genotypes at each developmental stage. Log₂-fold-changes were shrunk with an adaptive-shrinkage prior (ashr v2.2-54) and genes with Benjamini–Hochberg FDR < 0.05 were deemed significant. Time-dependent genes were detected via a likelihood-ratio test (LRT) contrasting full (genotype + developmental stage) against reduced (genotype) models. Pathway & gene-set enrichment: single-sample enrichment scores were computed from vst matrices using ssGSEA (GSVA v1.48.0) for MSigDB Hallmark (H) and KEGG (C2) collections obtained via msigdbr v7.5.1. Scores were analyzed with limma v3.56.2 using the same contrast matrix, with FDR < 0.05 marking significance. Complementary preranked GSEA (clusterProfiler v4.6.2) used signed log₂-fold-change lists, 10 000 phenotype permutations, and gene-set sizes between 10 and 500. The plots were rendered with ggplot2 v3.5.0, heatmap v1.0.12, EnhancedVolcano v1.18.0 and UpSetR v1.4.0.

#### scRNA-seq and analysis of small intestine lamina propria

Leukocytes were isolated from small intestine lamina propria (as detailed above) and flow-sorted using a gating strategy to isolate CD45^+^live cells. Cells were collected in low binding tubes containing PBS 0.05% (RNase-free BSA, Sigma, 126615-25ml). For each sample, an aliquot of cells was stained with acridine orange/propidium iodide Cell Viability Kit (Logos Biosystems, F23001) and counted with the LunaFx7 automatic cell counter (Logos Biosystems). Approximately [5000-30,000] cells were loaded on Chromium Chip and partitioned in nanolitre scale droplets using the Chromium X and Chromium GEM-X Single Cell Reagents (Chromium GEM-X Single Cell 3’ v4 Gene Expression User Guide, CG000731). Within each droplet the cells were lysed, and the RNA was reverse transcribed. All of the resulting cDNA within a droplet shared the same cell barcode. Illumina compatible libraries were generated from the cDNA using Chromium GEM-X Single Cell library reagents in accordance with the manufacturer’s instructions (Chromium GEM-X Single Cell 3’ v4 Gene Expression User Guide, CG000731). Final libraries are QC’d using the Agilent TapeStation and sequenced using the Illumina NovaSeq X. Sequencing read configuration: 28-10-10-90.

FASTQ files were aligned to the mm10 (GRCM38, version 93) transcriptome, and count matrices were generated, filtering for GEM cell barcodes (excluding GEMs with free-floating mRNA from lysed or dead cells) using Cell Ranger (version 9.0.0). All processing beyond this point was carried out using R version 4.4.1 using Seurat (version 5.2.1). Cells which did not meet the following criteria were removed; mitochondrial content within three standard deviations from the median, less than 500 genes detected per cell, and less than 1,000 RNA molecules detected per cell. Doublets were identified using DoubletFinder (version 2.0.4) and scDblFinder (version 1.20.2); cells called doublets by both methods were removed. Samples (4 biological replicates per Control and C5^ΔVav^ genotypes) were integrated using the RunHarmony function, using 3000 variable genes and the pca reduction on the first 20 principal components; as determined using the intrinsicDimensions (version 1.2) package; to construct the UMAP. Clusters were identified using a range of resolutions generating between 8 and 18 clusters. Clusters were annotated using a range of annotation packages at both the cluster and cell level : scCATCH (version 3.2.2); SingleR (version 2.8.0); clustifyr (version 1.18.0); scMCA (version 0.2.0) and cellTypist (version 1.6.3) and scanpy (version 1.10.4) within python (version 3.12.8). After inspection of marker genes and automated annotation results, a total of 12 clusters were identified using a resolution of 0.4. Marker genes per cluster were determined using the FindAllMarkers function, using a Wilcoxon rank-sum test, comparing each cluster to all other clusters. Differential gene expression between C5^ΔVav^ mutants and Control samples per cluster was determined using the GlmGamPoi package, using aggregated expression after pseudobulking, taking into account the 4 replicates per genotype. Gene Set enrichment analysis using the differential genelist was done using the fgsea (version 1.32.2) package with pathway and biological processes genesets download from Broad Institute, ranking the genes using the log2FC. Genesets were deemed statistically significant if their adjusted p.value < 0.05. Cell-cell communication between clusters was determined using CellChat (version 2.1.2). CellChat was run individually for the Control and C5^ΔVav^ samples, and major signalling changes across different Genotypes compared using quantitate contrasts and joint manifold learning, using the multiple datasets workflow. The cellchat analysis was performed as outlines in the cellchat vignette, with the population.size parameter set as ‘TRUE’, when computing the communication probability between clusters.

#### Ex vivo polyclonal activation of T cells

Dissociated mesenteric lymph nodes (mLNs) were isolated from C5^HetVav^ or C5^ΔVav^ were filtered through a 100μm cell strainer and counted. Lymphocytes were activated with CD3/CD28 coated beads following the manufacturer’s instructions (Miltenyi Biotec-130-093-627). 10^6^ mLN cells were incubated with 1:1 anti-CD3/CD28 coated beads for 24h at 37°C, 5% CO_2_ in 12 well plates (1mL total volume) with RPMI supplemeted with 10% fetal bovine serum (Gibco, 10270-106), L-glutamine (Stem Cell technologies, 07100), penicilin/streptramycin (Gibco, 15140-122), non-essential amino acids (Gibco, 11140-035), and sodium pyruvate (Gibco, 11360-039). After incubation, cells were stained with antibodies against CD3, CD4, CD8, CD69, CD44, and considered activated according to CD69 and CD44 expression in comparison to inactivated controls.

#### Lymphocyte transfer model of intestinal inflammation

Splenic lymphocytes were obtained from heterozygotes or homozygotes *Arpc5*^Cre-Foxp3-YFP^ mice and CD45^rbhi^ or Foxp3-YFP positive cells were FACS sorted and transferred into Rag^−/−^ mice. Positive control mice only received 5×10^5^ CD45^rbhi^ cells while adoptive transfer of 2×10^5^ Foxp3-YFP cells were given in the other groups as indicated in the figure. Mice weight was monitored and humane endpoint observed when reaching a 20% loss from starting weight.

#### Differentiation of bone marrow derived macrophages

Bone marrow was collected from the tibia and femur of male and female mice at 6 – 10 weeks and differentiated in vitro for 7 days in RPMI (Gibco R8758) supplemeted with 10% fetal bovine serum (Gibco, 10270-106), 10% L929 conditioned medium (Crick Cell services STP), L-glutamine (Stem Cell technologies, 07100), penicilin/streptramycin (Gibco, 15140-122), non-essential amino acids (Gibco, 11140-035), and sodium pyruvate (Gibco, 11360-039) to obtain bone marrow derived macrophages (BMDM).

#### Phagocytosis, bacterial killing and cytotoxicity assays in BMDM

To assess bacterial phagocytosis, adhered BMDMs (5 x 10^5^) were incubated with FITC-Heat-killed (HK) *E.coli* (60μg/mL-E2861-Invitrogen) or pHrodo *E.coli* (1μg/mL-P35366-Invitrogen) for up to 120 min at 37°C, 5% CO_2_. Negative controls were incubated at 4°C during the same period. Bacterial uptake was assessed by flow cytometry using the phagocytosis index calculation as follows: [(MFI x % positive cells at 37°C) – (MFI x % positive cells at 4°C)].

For bacterial killing (3×10^5^ BMDMs in 12 well plates) and cytotoxicity assays (10^5^ BMDMs in 96 well plate) adherent BMDM were incubated with phase *E. coli* K-12 (NCTC_10538) mid-log at MOI 5, 10 or 50. The bacteria concentration was assessed using OD_600_. The BMDM killing capacity was assessed by microscopy. Ten random fields of view were imaged on a Zeiss Axio Observer spinning-disk microscope equipped with a Plan Achromat 63×/1.4 Ph3 M27 oil lens, an Evolve 512 camera, and a Yokagawa CSUX spinning disk. The microscope was controlled by the SlideBook software (3i Intelligent Imaging Innovations). Images were analyzed using FIJI and internalized bacteria were detected using anti-E coli (abcam, ab137967). Microscopy secondary antibodies (A31572, Thermo Scientific) and DAPI (4083S, Cell Signalling Technology). Calculations were obtained as follows: [(internalized bacteria at T0) – (internalized bacteria at Time X)].

The BMDMs cytotoxicity was assessed using the CyQUANT^Tm^LDH Cytotoxicity assay following the manufacturer’s instructions (Thermo Fisher Scientific, C20300). For G-actin or F-actin detection, BMDMs (5×10^5^ in non-adherent 6 well plates) were incubated with E. coli K-12 (MOI 50) for 15 min or vehicle. 30 min pre-treatment with 10µM Cytochalasin D (Merck, 250255) was performed when indicated. After treatment cells were permeabilized using the Foxp3/Transcription Factor Staining Buffer Set (00-5523-00 ThermoFisher Scientific) and stained with antiDNAse-I Alexa Fluor 488 (Invitrogen, D12371) or Alexa Fluor 568-phalloidin (Molecular Probes), respectively.

#### Morphological and live cell imaging of BMDM

BMDM^LifeAct-GFP^ derived from C5^HetVav^ or C5^ΔVav^ murine bone marrow were seeded on fibronectin coated 35 mm Matek dish at 5×10^4^ cells/well and incubated overnight at 37°C with 5% CO_2_. Cells were imaged as described above. Data for 10-12 cells per sample were segmented using ilastik and segmented images were then analyzed using FIJI. For live cell imaging the cells were seeded on fibronectin coated ibidi 4-well glass-bottom µ-slide at 4×10^4^ cells/well and incubated overnight at 37°C with 5% CO_2_. pHrodo™ Deep Red E. coli BioParticles™ Conjugate (Thermo Fisher) was prepared according to manufacturer’s instructions. Prior to imaging, 5 µL of pHrodo™ Deep Red E. coli was added into each well. The imaging chamber was maintained at 37°C. Both 2D and 3D live cell imaging of phagocytosis used a Zeiss Axio Observe microscope, an Evolve 512 camera and a Yokagawa CSUX spinning disk. For 2D live cell imaging, cells were imaged at a single z plane every 500 ms for 3 minutes using a Plan Achromat 100x/1.46 lens. For 3D live cell imaging, cells were images every 10 s for 4 minutes, with Z-stack of 0.27 µm steps over 10 µm with a Plan Achromat5 63x/1.4 Ph3 M27 oil lens. The microscope was controlled by the SlideBook software (3i Intelligent Imaging Innovations). The 2D videos were generated using Fiji and the 3D videos were reconstructed using napari.

#### Statistical analysis

Data are shown as the mean ± S.E.M. Sample sizes were established prior to experimental approach with help of the experimental design assistant of NC3Rs to achieve statistical power while minimizing animal use. To test normality or lognormality we applied D’Agostino & Pearson test. All statistical comparisons were performed using Prism 10 (GraphPad) with specific statistical test applied as stated in the figure legends. Statistical significance was considered as *P*<0.05.

**Suppl.1. (A)** Arpc5 deletion strategy using Vav1-iCre and the level of Arpc5 and Arpc5l in C5^fl/fl^, C5^HetVav^ or C5^ΔVav^ bone marrow extracts. (**B**) Weight change (g) of C5^fl/fl^, C5^HetVav^ or C5^ΔVav^ mice over 15 weeks. (**C**) Representative H&E images of the ileum rolls. (**D**) Macrophages and neutrophils quantification in different intestinal sections (**E**) Quantification of the histopathology score of intestines of indicated 8-15 wks old mice. (**F**) Cell numbers of mesenteric lymph nodes (mLNs) B and T cells, CD4^+^ and CD8^+^, macrophages (MØs), neutrophils (NØs) and dendritic cells (DCs) in the indicated mice. (**G**) The levels of Arpc5 and Arpc5l in C5L^HetVav^ or C5L^ΔVav^ bone marrow extracts, actin is a loading control. (**H**) Representative H&E images of ileum rolls and Lipocalin-2 (LPCN-2) faecal levels in the indicated mice. (**I**) Cell numbers of B, CD4+, CD8^+^ and natural killer (NK) lymphocytes in the blood of indicated mice. (**J**). NØs, MØs, DCs, B lymphocytes, CD4^+^, CD8^+^ and NK cells numbers in the spleen of C5^HetVav^ or C5^ΔVav^ mice. (**K**) Representative H&E images and pathological score of ileum rolls of indicated animals after bone marrow transplantation. Quantification of the levels of C-reactive protein (CRP) in the serum, LPCN-2 in the faeces and spleen weight in indicated animals after bone marrow transplantation. Data shown as means ± SEM of at least three independent experiments, n=8. Statistical analysis was performed using two-way analysis of variance (ANOVA) in **B**; Kruskal-Wallis test in **D**, **E, F**, **H, I** and **J**; Mann-Whitney test in **K** and significant *p* values are indicated on the graphs. C5^fl/fl^ (Arpc5^fl/fl^), C5^HetVav^ (Arpc5^HetVav^), C5^ΔVav^ (Arpc5^ΔVav^) and C5L^ΔVav^ (Arpc5l^ΔVav^). Scale bars = 500µm. ns = not significant. * = p-value < 0.05. ** = p-value < 0.01. *** = p-value < 0.001 and **** = p-value < 0.0001.

**Suppl.2.** (**A**) Representative H&E image of ileum rolls of 4 week old (wk) C5^HetVav^ or C5^ΔVav^ mice. (**B**) Number of differentially expressed gene (DEG) pathways (left) and enriched DEG hallmark (right) observed in intestinal lamina propria macrophages obtained from C5^HetVav^ or C5^ΔVav^ mice over the time. (**C**) Total cell numbers in mLNs of 4 and 8 week old (left) together with the cell counts of B, T and CD4^+^ or CD8^+^ lymphocytes (middle), neutrophils (NØs), macrophages (MØs) or dendritic cells (DCs) (right) in mLNs from 4 week (wk) old C5^HetVav^ or C5^ΔVav^ mice. (**D**) Splenic B, T and natural killer (NK) lymphocytes (left), NØs, MØs and DCs (right) cell counts in the indicated mice after antibiotic treatment. (**E**) mLNs total cell counts (left), together with NØs and MØs (middle), B and T lymphocytes (right) cell numbers from mLNs of mice treated with antibiotic. (**F**) Prevalence fraction of most abundant microbial classes found in indicated mice intestines. Data shown as means ± SEM of at least three independent experiments, n=7, except in **F** where n=8. Statistical analysis was performed using Kruskal-Wallis test in **C** left, **D** and **E**; Multiple Mann-Whitney test was performed in **C** right. Multiple Mann-Whitney test was performed in **F**. Significant *p* values are indicated on the graphs. Controls used were C5^HetVav^ (Arpc5^HetVav^), C5^ΔVav^ (Arpc5^ΔVav^), Veh-C5^ΔVav^ (Arpc5^ΔVav^ treated with vehicle), Atbx-C5^ΔVav^ (Arpc5^ΔVav^ treated with antibiotics). Scale bars = 500µm. ns = not significant. * = p-value < 0.05. ** = p-value < 0.01. *** = p-value < 0.001 and **** = p-value < 0.0001.

**Suppl.3.** (**A**) Bar plots representing the proportion of immune cells per biological replicate in C5^HetVav^ or C5^ΔVav^ mice. (**B**) Biological relevant pathways of differentially expressed genes in Macrophages, Monocytes and Neutrophils from C5^HetVav^ and C5^ΔVav^ animals. (**C**) Anti CD3/CD28 polyclonal activation assay in T cells from C5^HetVav^ (Arpc5^HetVav^) and C5^ΔVav^ (Arpc5^ΔVav^) animals. (**D**) Representation of the differential interaction strength obtained from CellChat analysis. (**E**) Expression levels of IL-10 and IL-10 receptor (IL-10R) (left). IL-10 signalling communication with outgoing (IL-10 production) and incoming (IL-10R) interactions between T regulatory cells (Tregs), Macrophages and Monocytes inferred using CellChat (right). (**F**) Tregs function in the lymphocyte transfer model of intestinal inflammation. CD45^rbhi^, CD45^rbhi^ + Foxp3^WT^ or CD45^rbhi^ + Foxp3^C5KO^ were transplanted into Rag^−/−^ mice and weight change and survival probability assessed over four weeks. Mann-Whitney test was performed in **C**. ns = not significant.

**Suppl.4.** (**A**) Spleen weight, faecal lipocalin-2 (LPCN-2) levels (left) and representative H&E images of cecum of control^RagKO^ or C5^ΔVavRagKO^ mice (right). (**B**) Most abundant classes of bacteria found in the intestinal microbiome analysis of control^RagKO^ or C5^ΔVavRagKO^. (**C**) Representative Facs plots (left) and quantification of CD45.1^+^ or CD45.2^+^ of mice after bone marrow transplantation (right). (**D**) Spleen weight (left) and faecal lipocalin-2 (LPCN-2) levels (right) in indicated mice after bone marrow transplantation. (**E**) Representative image of H&E images of ileum rolls. (**F**) Quantification of the levels of serum C-reactive protein (CRP), spleen weight, mLNs cell counts and immature neutrophils (NØs) counts in the bone marrow of indicated animals after busulfan adoptive transfer. (**G**) CRP levels in the serum and spleen weight of indicated animals after busulfan depletion followed by mononuclear phagocytes (MNPs) adoptive cell transfer. (**H**) CRP levels in the serum, LPCN-2 faecal levels and histopathology score together with representative H&E images of the ileum of clodronate liposomes depleted 4 week old animals followed by wild-type (MNP^WT^) or C5^ΔVav^ (MNP^KO^) transplantation. Data shown as means ± SEM of at least three independent experiments, n=8, except in B and G where n=5 from two independent experiments. Statistical analysis was performed using Mann-Whitney test in **A**, **B**, **D** and **H;** Kruskal-Wallis test in **F and G**. Significant *p* values are indicated on the graphs. Controls used were Arpc5^HetVav^, C5^ΔVav^ (Arpc5^ΔVav^). Scale bars are indicated in the figure. ns = not significant. * = p-value < 0.05. ** = p-value < 0.01. *** = p-value < 0.001 and **** = p-value < 0.0001.

**Suppl.5.** (**A**) Representative Facs plot and measurements of side scatter area (SSC-A) or forward scatter area (FSC-A) of indicated bone marrow derived macrophages (BMDMs). (**B**) Facs histograms of G-actin or F-actin in unstimulated BMDMs. (**C**) Quantification of *E coli*-FITC uptake by C5^+/−^ or C5^−/−^ BMDMs in the indicated timepoints. (**D**) Representative F-actin Facs histograms of *E.coli*-activated BMDMs. Data shown as means ± SEM of a representative of at least three independent experiments. Statistical analysis was performed using Mann-Whitney test in **A** and two-way ANOVA in **C**. Significant *p* values are indicated on the graphs. C5^+/−^ = Arpc5^+/−^, C5^−/−^ = Arpc5^−/−^. *** = p-value < 0.001 and **** = p-value < 0.0001.

**Movie 1.** Phagocytosis of E.coli by BMDMs with and without Arpc5. Representative movie showing actin dynamics in Arpc5^+/−^ (left) and Arpc5^−/−^ (right) BMDMs expressing GFP-LifeAct (Cyan) during phagocytosis of *E. coli* (Red). White arrows indicate the bacteria being phagocytosed. The time in minutes and seconds is shown on the upper left corner. Scale bar = 10 µm. Image stills from the movie are shown in Fig. 4D.

**Movie 2.** 3D view of *E. coli* uptake by BMDMs. Representative movie showing *E. coli* (Red) uptake in BMDMs expressing GFP-LifeAct (Cyan) in Arpc5^+/−^ (left) and Arpc5^−/−^ (right) BMDM. The time in minutes is shown on the upper left corner and the scale bar is shown on the lower right corner. Image stills from the movie are shown in Fig. 4E.

## Notes

### Competing Interest Statement

The authors have declared no competing interest.

### Summary of Updates

New data has been added to address reviewer questions and a new author added.

