## Supplementary figures and images for "Branched actin networks mediate macrophage-dependent host microbiota homeostasis"

### Combined revised supplemental figures

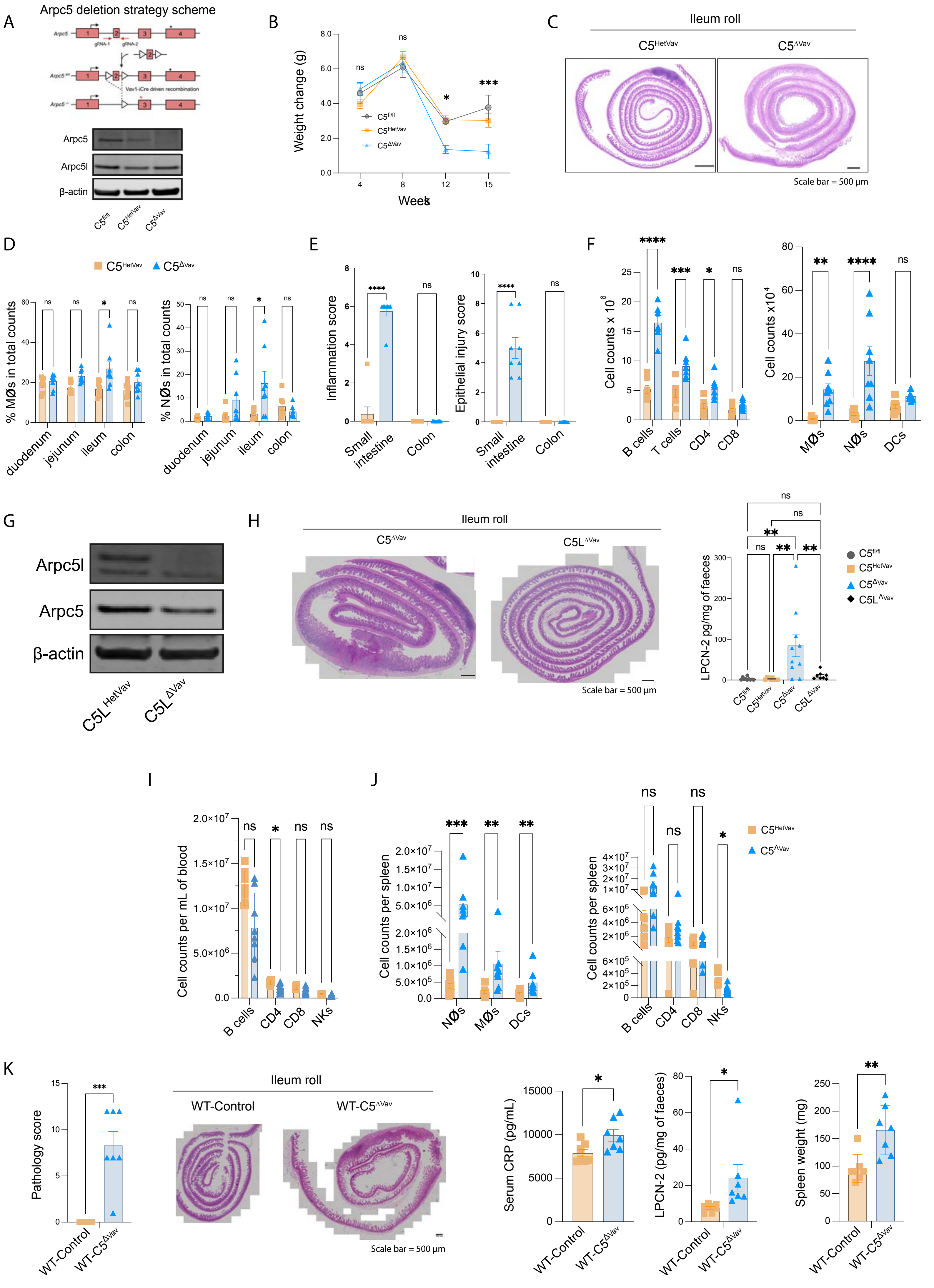

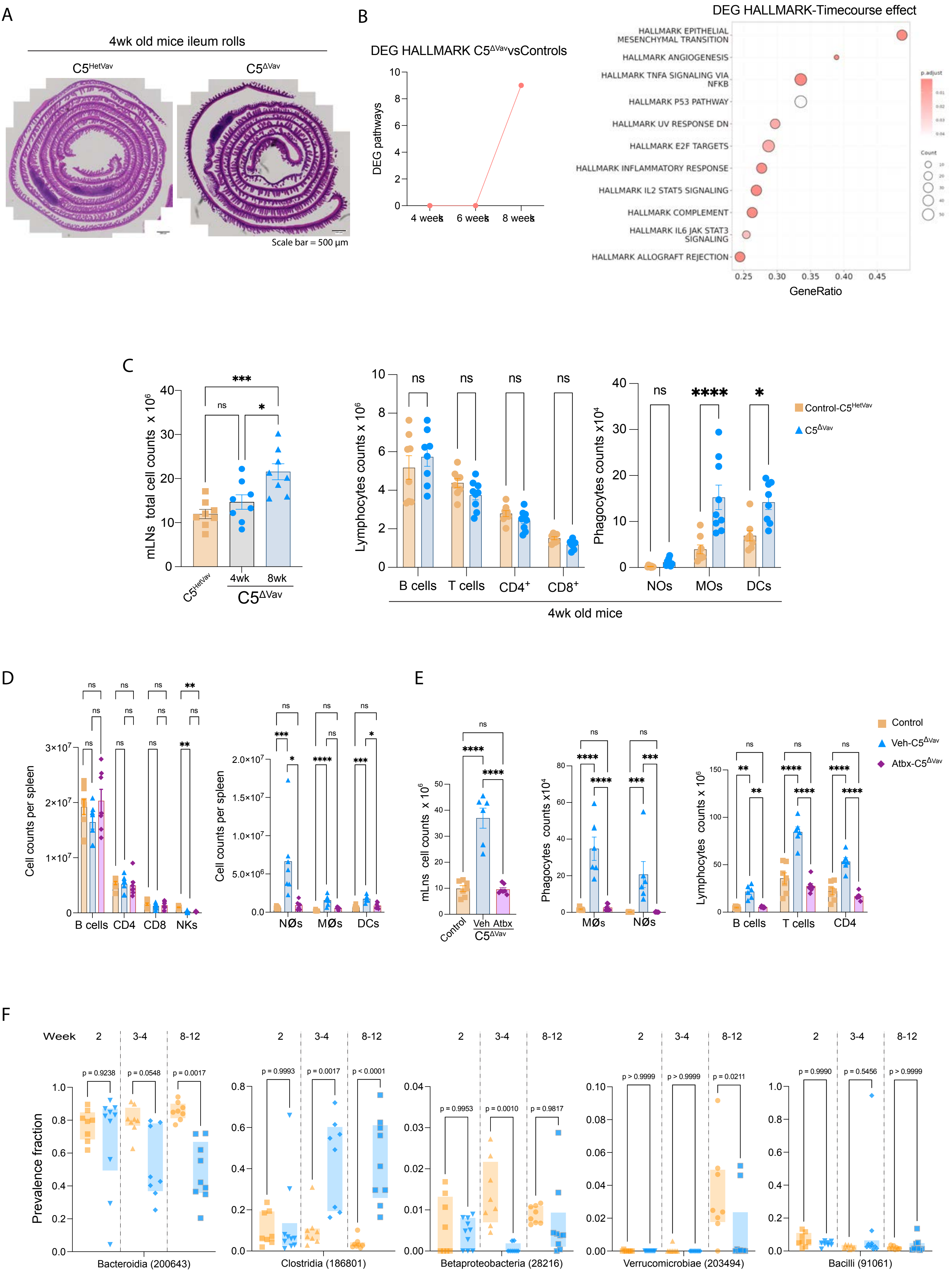

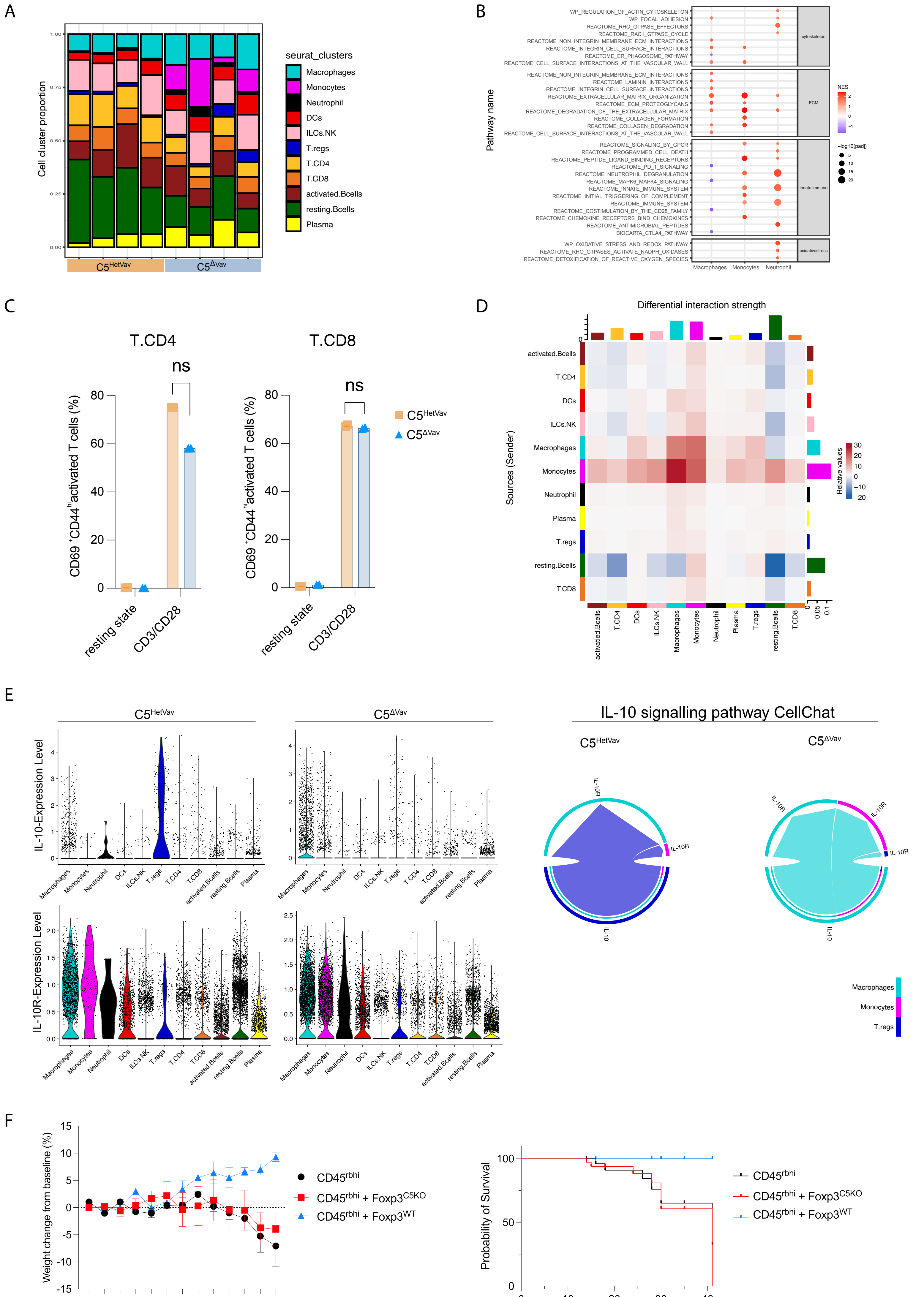

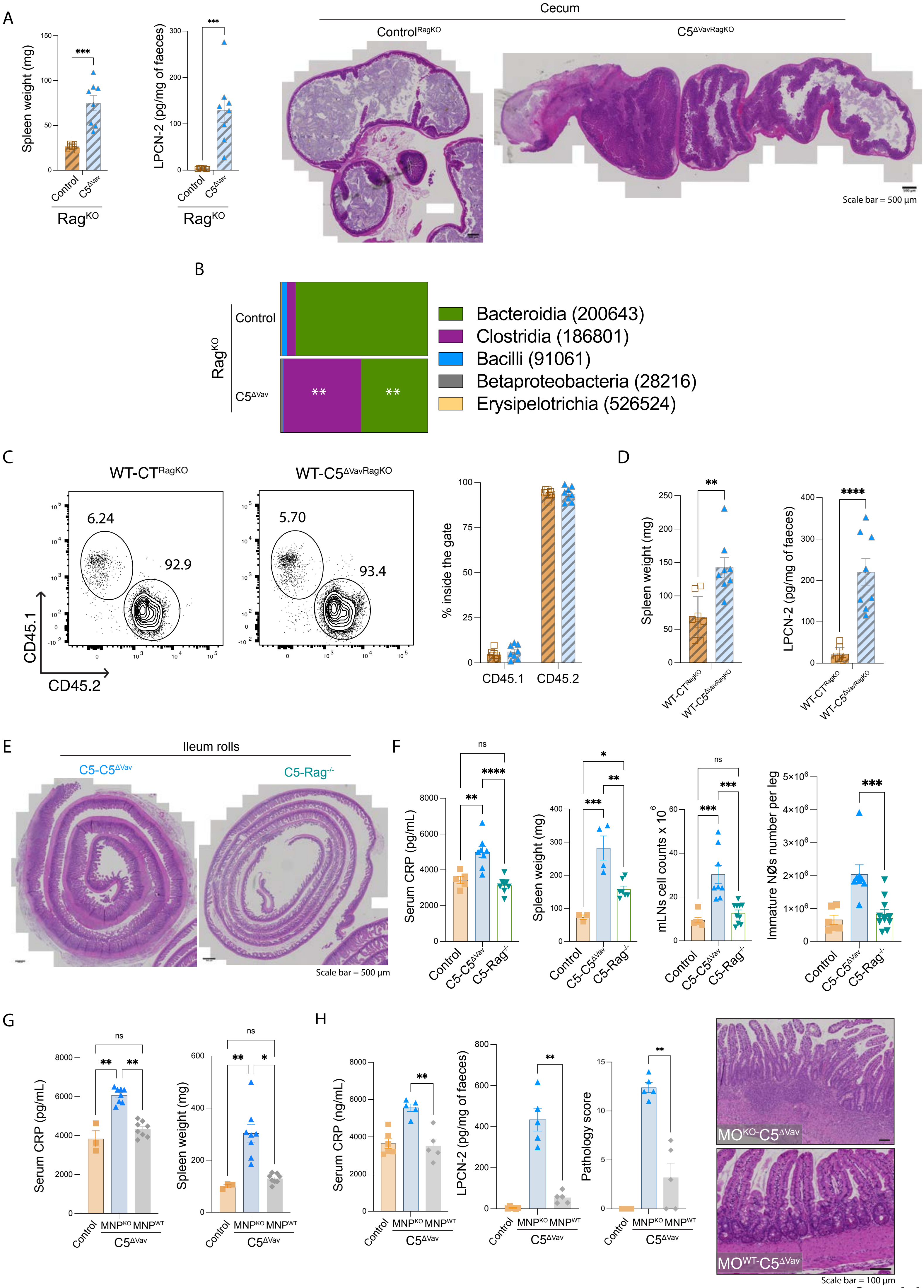

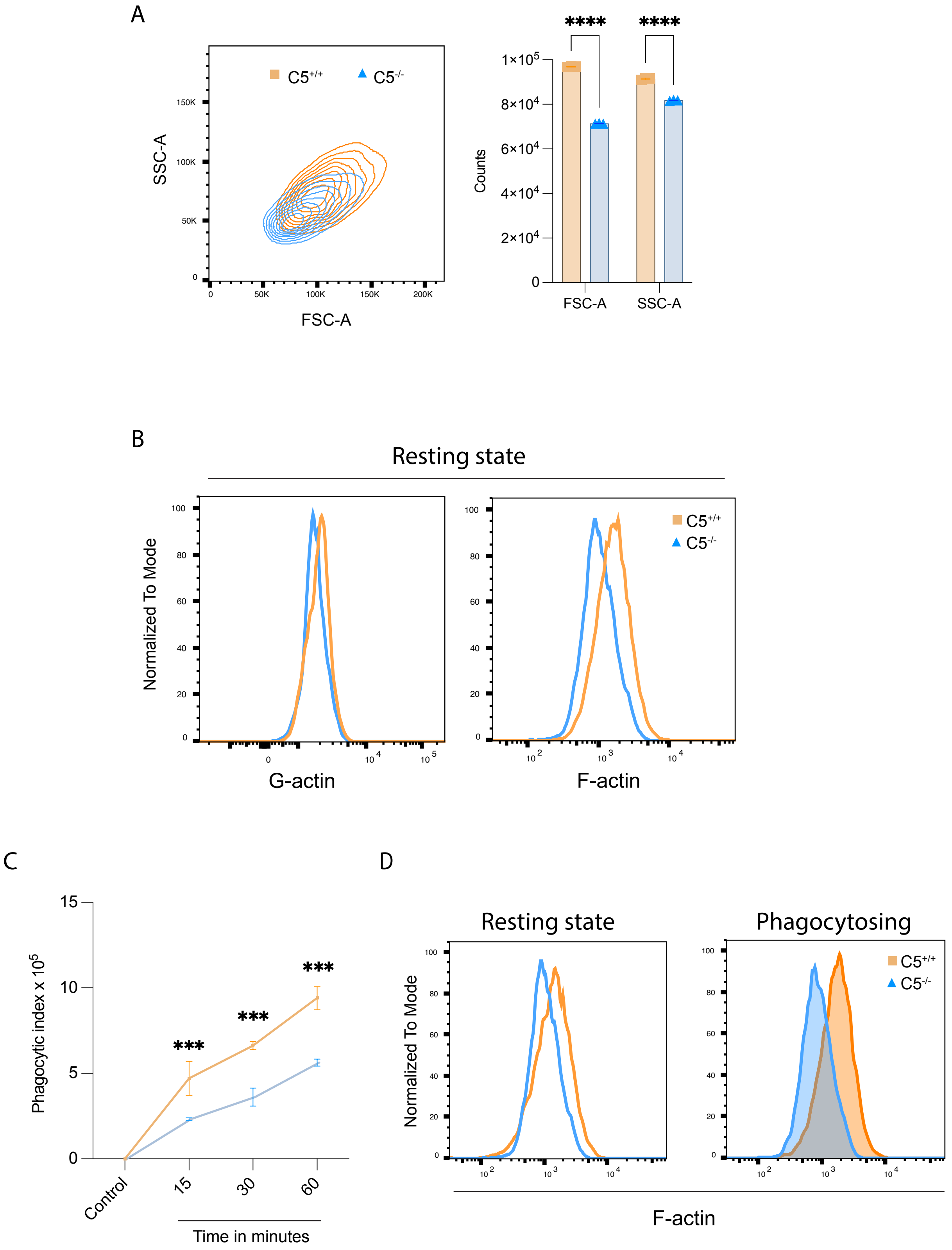
